## Supplemental Figures for "Comprehension of computer code relies primarily on domain-general executive brain regions"

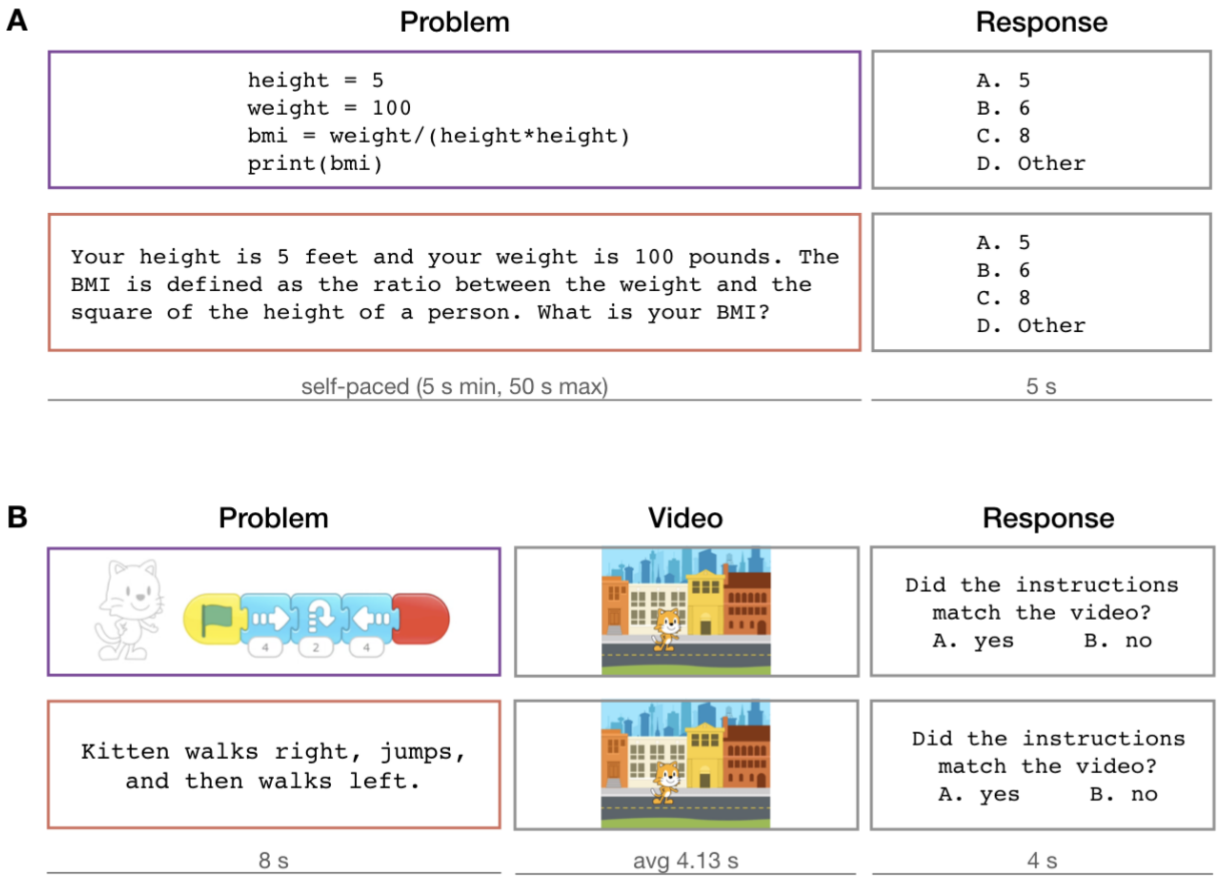


***[Figure 1 – supplement 1]****. Trial structure of the critical task. (A) Experiment 1 - Python (B) Experiment 2 - ScratchJr. All analyses use fMRI responses to the "problem" step.*

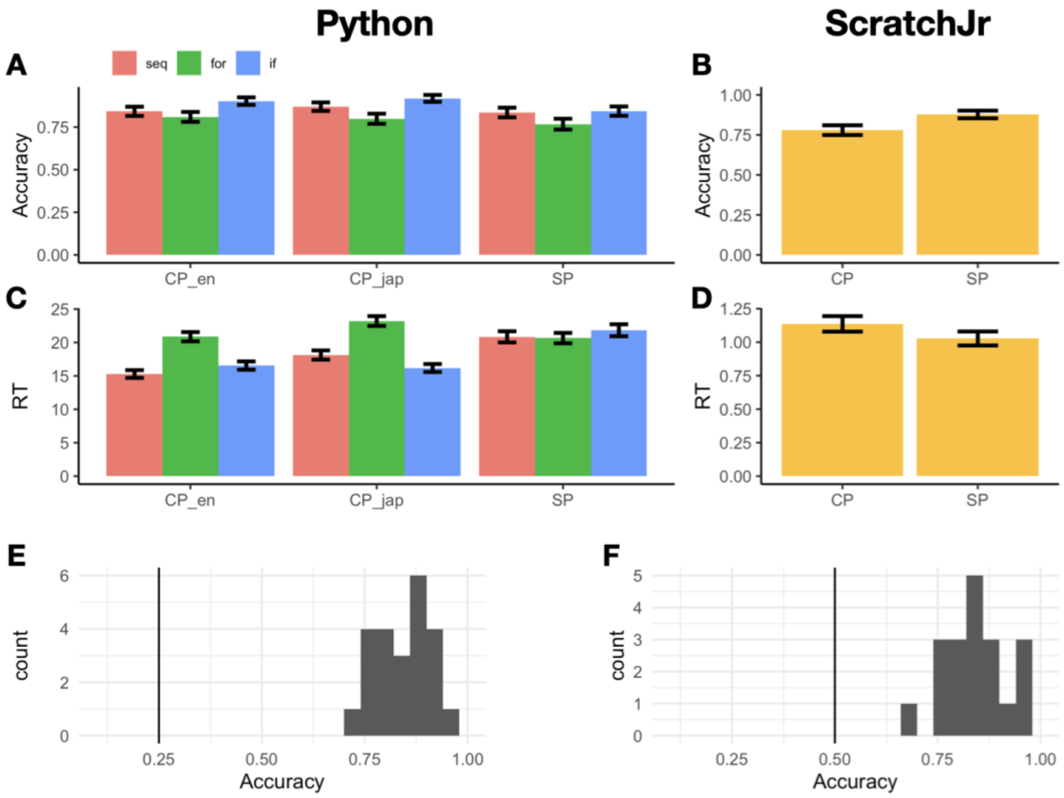


**[Figure 2 – supplement 1].** *Behavioral results. (**A**) Python code problems had mean accuracies of 85.1% and 86.2% for the English-identifier (CP_en) and Japanese-identifier (CP_jap) conditions, respectively, and sentence problems (SP) had a mean accuracy of 81.5%. There was no main effect of condition (CP_en, CP_jap, SP), problem structure (seq – sequential, for – for loops, if – if statements), or problem content (math vs. string); however, there was a three-way interaction among Condition (sentence problems > code with English identifiers), Problem Type (string > math), and Problem Structure (for loop > sequential; p = 0.02). Accuracy data from one participant had to be excluded due to a bug in the script. (**B**) ScratchJr code problems had a mean accuracy of 78.0%, and sentence problems had a mean accuracy of 87.8% (the difference was significant: p = 0.006). (**C**) Python problems with English identifiers had a mean response time (RT) of 17.56 s (SD = 9.05), Python problems with Japanese identifiers had a mean RT of 19.39 s (SD = 10.1), and sentence problems had a mean RT of 21.32 s (SD = 11.6). Problems with Japanese identifiers took longer to answer than problems with English identifiers (β*

*= 3.10, p = 0.002), and so did sentence problems (β = 6.12, p < 0.001). There was also an interaction between Condition (sentence problems > code with English identifiers) and Program Structure (for > seq; β = -5.25, p < 0.001), as well as between Condition (CP_jap > CP_en) and Program Structure (if > seq; β = -2.83, p = 0.04). There was no significant difference in response times between math and string manipulation problems. (**D**) ScratchJr code problems had a mean RT of 1.14 s (SD = 0.86), and sentence problems had a mean RT of 1.03 s (SD = 0.78; the difference was not significant. The RTs are reported with respect to video offset. Items where >50% participants chose the incorrect answer for the (easy) verbal condition were excluded from accuracy calculations. (**E**) Mean accuracies for all Python participants were above chance. (**F**) Mean accuracies for all ScratchJr participants were above chance.*

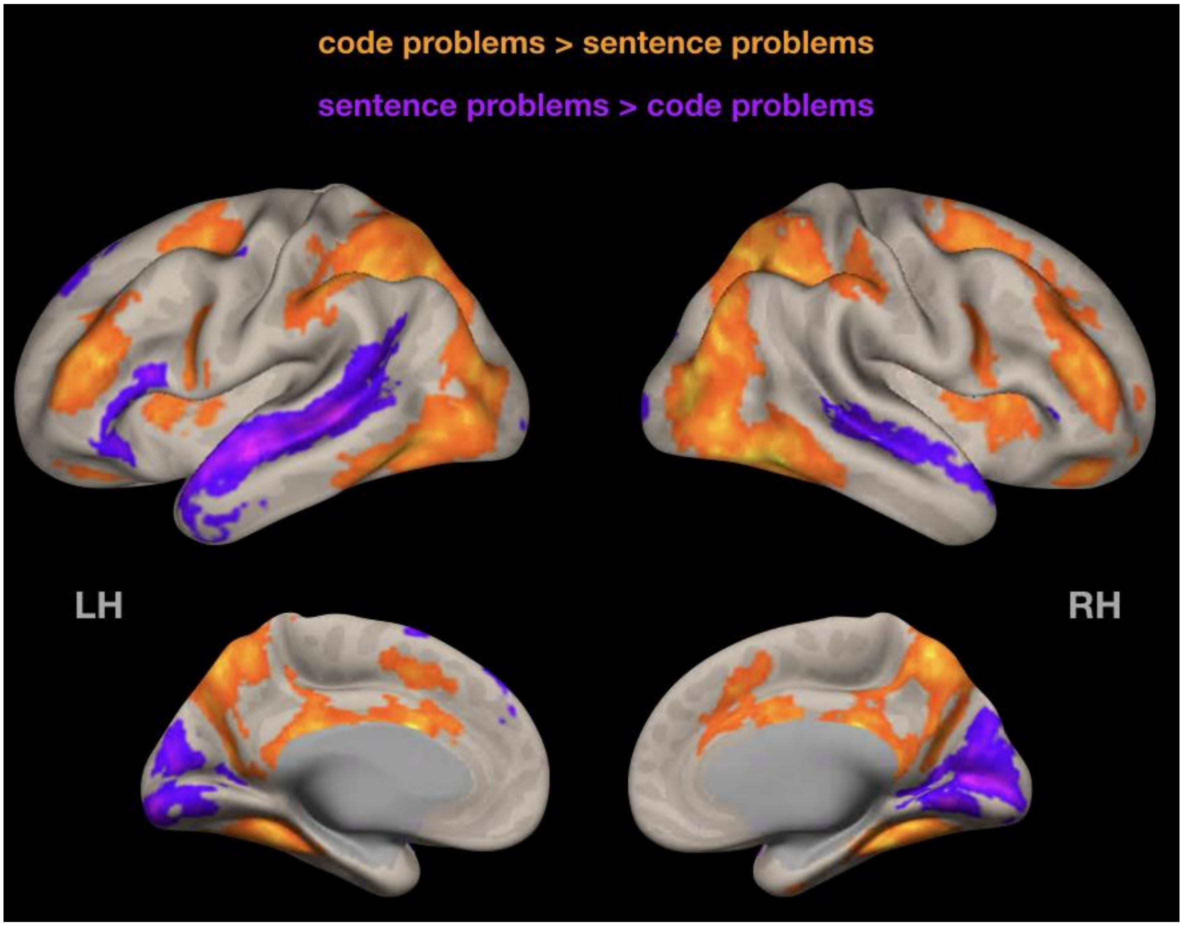


***[Figure 2 – supplement 2].*** *Random-effects group-level analysis of Experiment 1 data (Python, code problems > sentence problems contrast). Similarly to analyses reported in the main text, code-evoked activity is bilateral and recruits fronto-parietal but not temporal regions. Cluster threshold $p < 0.05$, cluster-size FDR-corrected; voxel threshold: $p < 0.001$, uncorrected.*

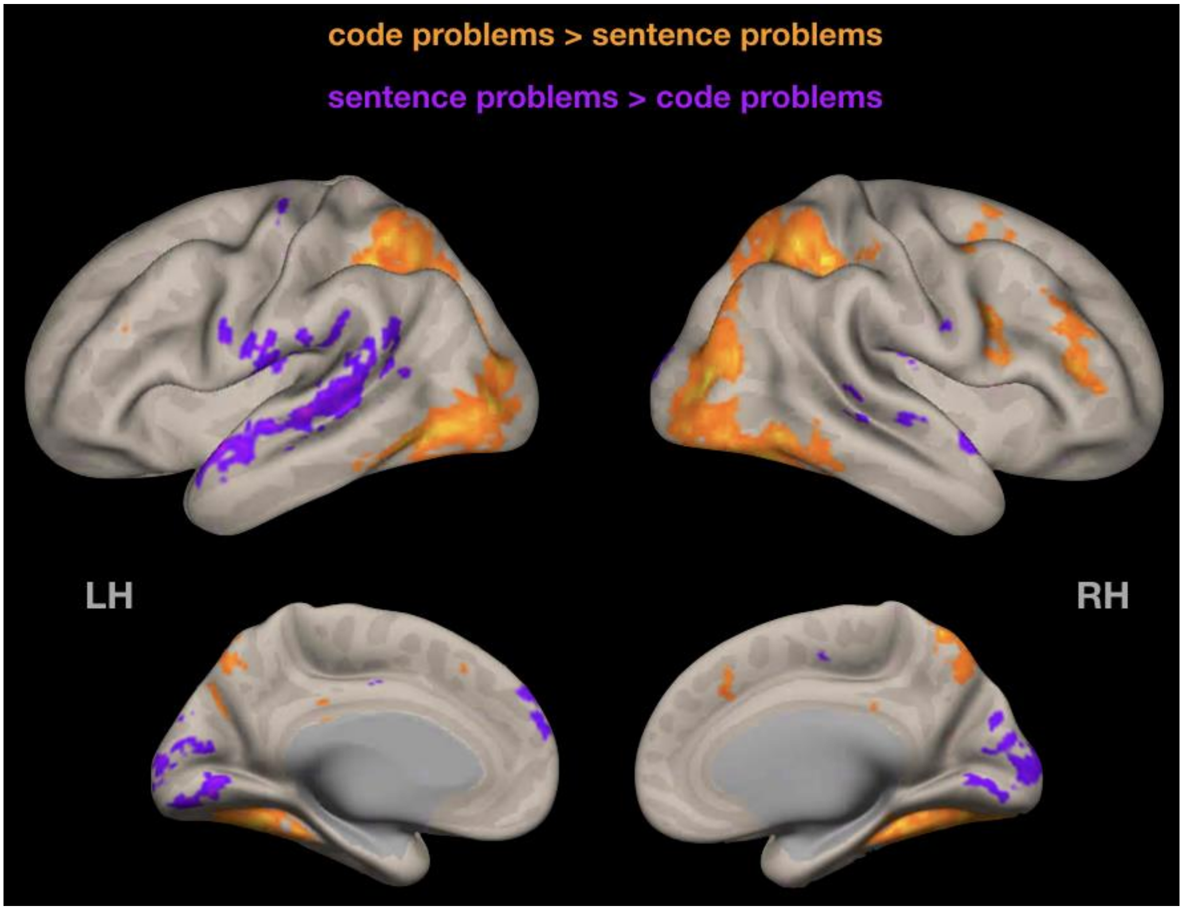


***[Figure 2 – supplement 3].*** *Random-effects group-level analysis of Experiment 2 data (ScratchJr, code problems > sentence problems contrast). Similarly to analyses reported in the main text, ScratchJr-evoked activity has a small right hemisphere bias. Cluster threshold $p < 0.05$, cluster-size FDR-corrected; voxel threshold: $p < 0.001$, uncorrected.*

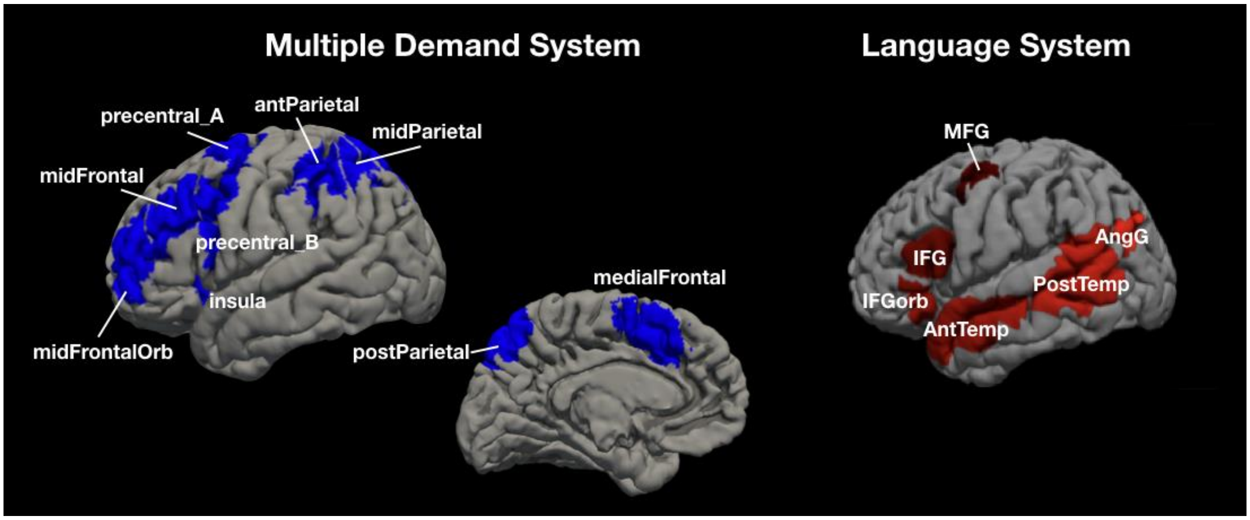


**[Figure 3 – supplement 1].** *The parcels in the two candidate brain systems of interest, multiple demand (MD) and language. The parcels are derived from group-level representations of MD and language activity and are used to define the functional regions of interest (fROIs) in individual participants (NB: we show the left hemisphere parcels for the MD system, but the system is bilateral). For each participant, the network of interest is comprised of the top 10% of voxels within each parcel with the highest t-value for the relevant contrast (MD - hard vs. easy spatial working memory task; language – sentence reading vs. nonword reading; see* ***Methods****). Abbreviations: mid – middle, ant – anterior, post – posterior, orb – orbital, MFG – middle frontal gyrus, IFG – inferior frontal gyrus, temp – temporal lobe, AngG – angular gyrus.*

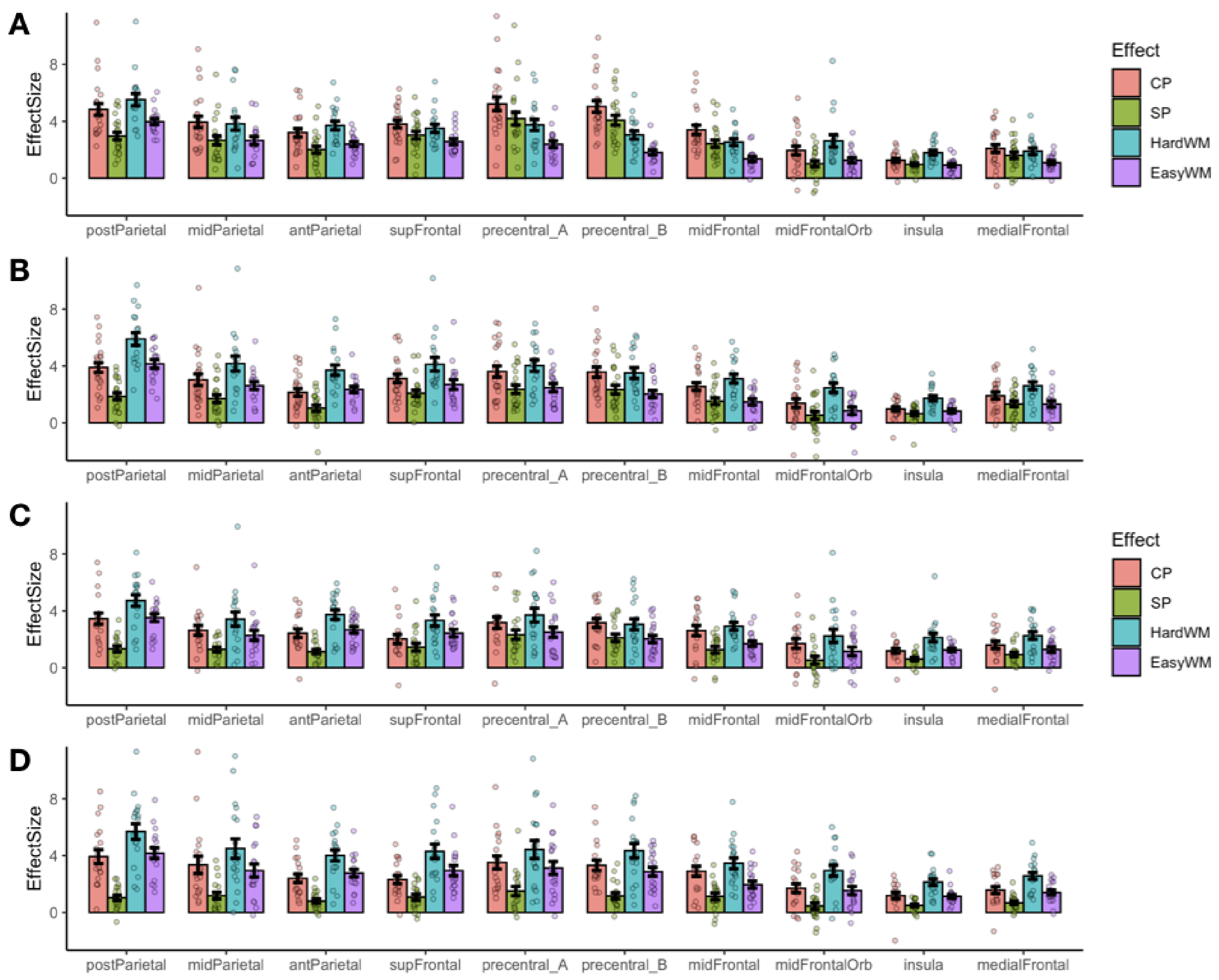


***[Figure 3 – supplement 2]****. ROI-level responses in the multiple demand system to the critical task (CP – code problems, SP – sentence problems) and the spatial working memory task (HardWM – hard working memory task, EasyWM – easy working memory task). (A) Experiment 1, Python; left hemisphere fROIs; (B) Experiment 1, Python; right hemisphere fROIs; (C) Experiment 2, ScratchJr; left hemisphere fROIs; (D) Experiment 2, ScratchJr; right hemisphere fROIs. No fROIs prefer both Python and ScratchJr code problems over the spatial working memory task.*

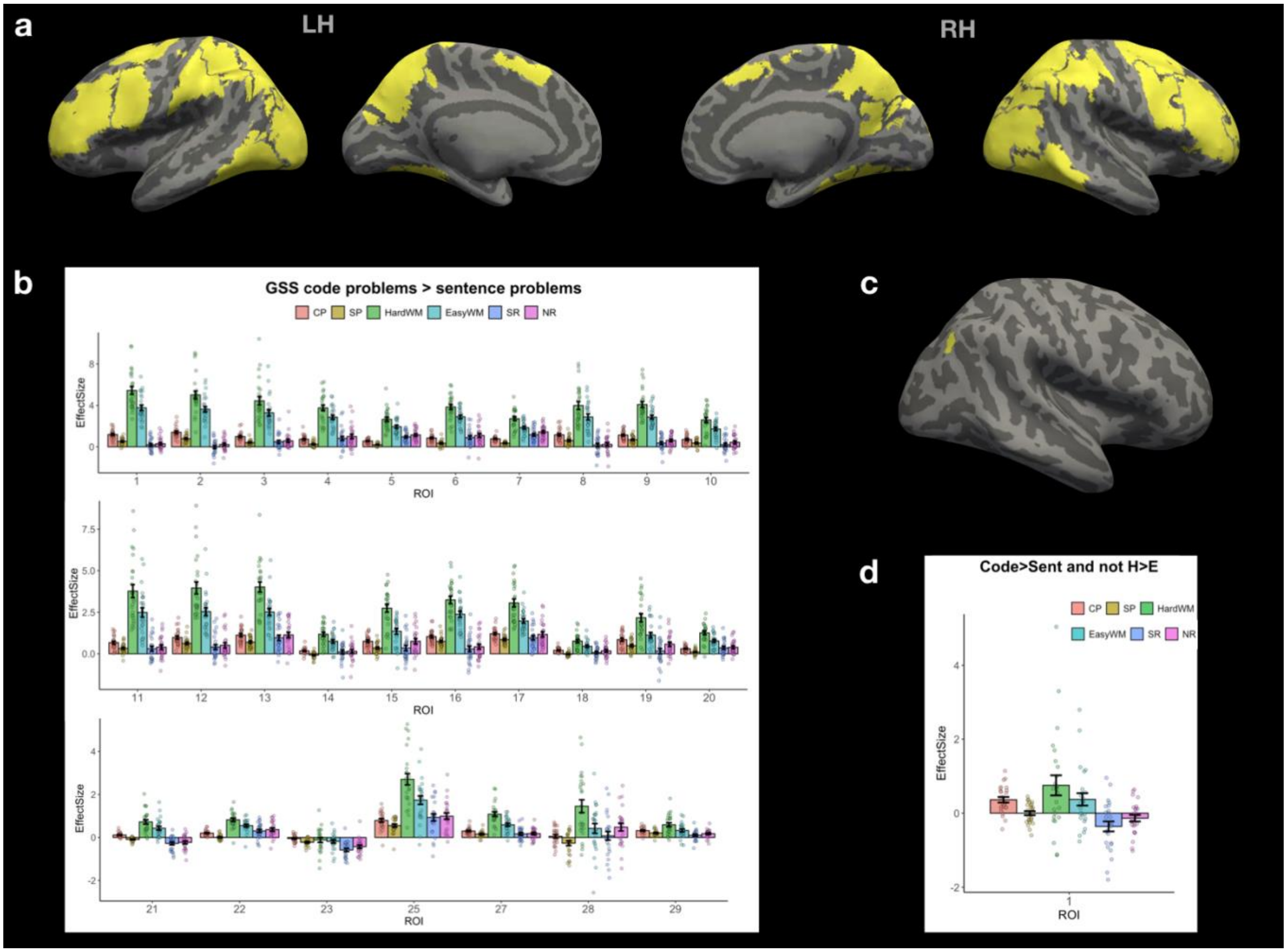


***[Figure 3 – supplement 3]***. *Whole-brain group-constrained subject-specific analysis (GSS; Fedorenko et al, 2010) based on data from Experiment 1 shows the absence of code-only brain regions. (a) Parcels defined on the group level using the code problems > sentence problems contrast, p threshold 0.001, inter-subject overlap >= 70%. (b) Activation profile for the top 10% of voxels within each parcel in (a) across conditions. All code-sensitive regions exhibit high activity during the spatial working-memory task, suggesting that they belong to the MD system. (c) Parcels defined using the contrast above plus the "not hard working-memory task > easy working-memory task" contrast, p=0.5. Only one parcel was significant (right hemisphere). (d) Even that parcel's response profile shows high activity in response to the working-memory task, modulated by difficulty, rather than a code-specific response. Abbreviations:*

*CP – code problems; SP – sentence problems; HardWM – hard working memory task; EasyWM – easy working memory task; SR – sentence reading; NR – nonword reading.*

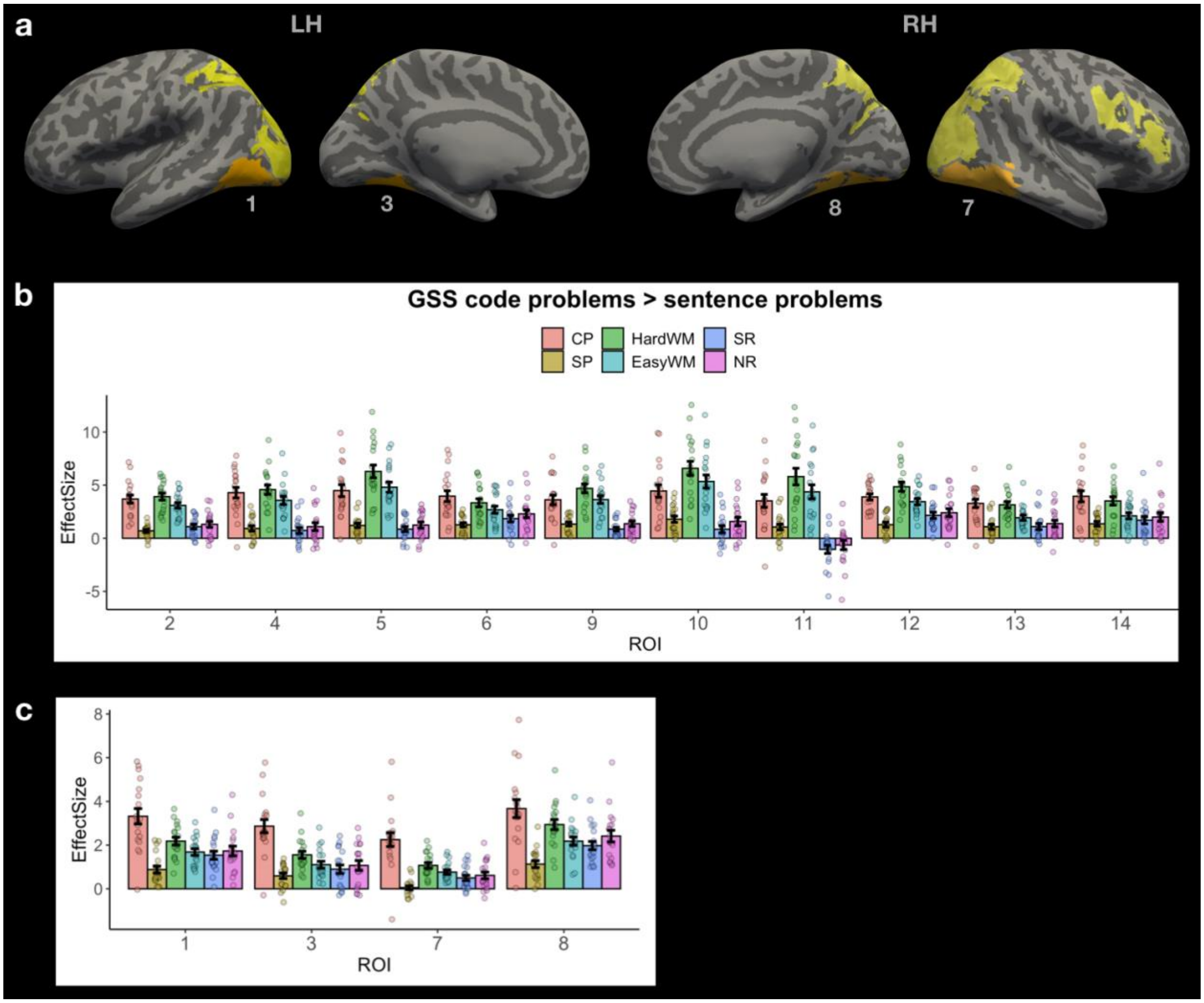


**[Figure 3 – supplement 4].** *Whole-brain group-constrained subject-specific analysis (GSS; Fedorenko et al, 2010) based on data from Experiment 2. (a) Parcels defined on the group level using the code problems > sentence problems contrast, p threshold 0.001, inter-subject overlap >= 70%. Parcels where the responses to ScratchJr code were stronger than responses to all other tasks are labeled and marked in orange; they include parts of early visual cortex and parts of the ventral visual stream. (b) Activation profile for the top 10% of voxels within each parcel in (a) marked in yellow. All regions exhibit high activity during the spatial working-memory task, suggesting that they belong to the MD system. (c)*

*Activation profile for the top 10% of voxels within each parcel in (a) marked in orange. These fROIs exhibit higher responses to ScratchJr problems compared to a working memory task; given that they are located in the visual cortex, we can infer that they respond to low-level visual properties of ScratchJr code. A follow-up conjunction analysis using the contrast in (a) plus the "not hard working-memory task > easy working-memory task" contrast, $p=0.5$, revealed no significant parcels, indicating the lack of code-selective response. Abbreviations: CP – code problems; SP – sentence problems; HardWM – hard working memory task; EasyWM – easy working memory task; SR – sentence reading; NR – nonword reading.*

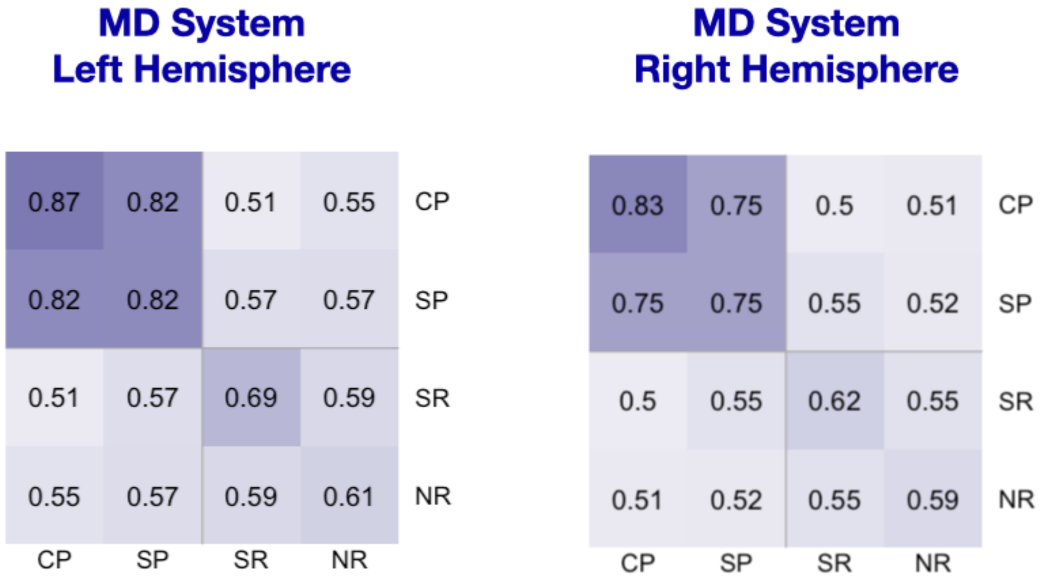


***[Figure 4 – supplement 1]****. Spatial correlation analysis of voxel responses within the MD system during the Python experiment (CP – code problems and SP – sentence problems) with the language localizer conditions for the same participants (SR – sentence reading and NR – nonword reading). Each cell shows a correlation between voxel-level activation patterns for each condition. Within-condition similarity is estimated by correlating activation patterns across independent runs. Code problems correlate with sentence problems much more strongly than with sentence reading ($\beta$ = -0.59, $p < 0.001$) and with nonword reading ($\beta$ = -0.55, $p < 0.001$), but substantially weaker than with other code problems ($\beta$ =*

*0.11, p < 0.001). There was no main effect of hemisphere, but there was an interaction between code/sentence problems and reading conditions (sentence reading: β = 0.17, p < 0.001, nonword reading: β = 0.13, p = 0.002), indicating that the correlation patterns of code/sentence problems were somewhat less robust in the right hemisphere.*

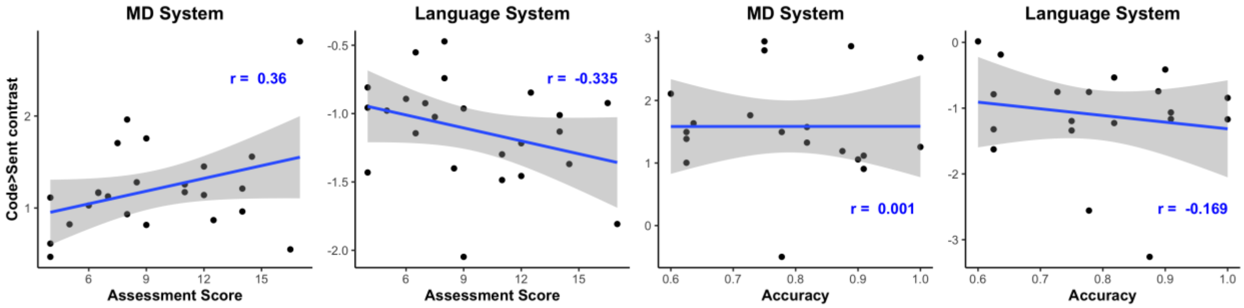


***[Figure 4 – supplement 2].*** *The effect of programming expertise on code-specific response strength within the MD and language system in Experiment 1, Python (**A, B**) and Experiment 2, ScratchJr (**C, D**). Python expertise was evaluated with a separate one-hour-long Python assessment (see the paper's website https://github.com/ALFA-group/neural-program-comprehension); ScratchJr expertise was estimated with in-scanner response accuracies. No correlations were significant.*
