## Supplemental File - fROI stats for "Comprehension of computer code relies primarily on domain-general executive brain regions"

### *Supplementary File – fROI stats*

***Table S1***. *Responses to Python code problems (CP) vs. sentence problems (SP) and nonword reading (NR) in individual fROIs within the multiple demand system. P values are Bonferroni-corrected for the number of regions. Non-significant values are italicized and marked in gray.*

| Hemisphere | fROI | Regression Term | Beta | p value |
|---|---|---|---|---|
| L | postParietal | Intercept | 4.79 | 2.00E-20 |
| | | CP>SP | 1.86 | 3.00E-07 |
| | | CP>NR | 4.17 | 1.05E-18 |
| L | midFrontal | Intercept | 3.39 | 1.56E-16 |
| | | CP>SP | 0.97 | 0.001 |
| | | CP>NR | 2.5 | 9.62E-14 |
| L | precentral_B | Intercept | 4.91 | 1.23E-14 |
| | | CP>SP | 0.95 | 0.002 |
| | | CP>NR | 3.38 | 1.68E-18 |
| L | antParietal | Intercept | 3.13 | 4.40E-15 |
| | | CP>SP | 1.17 | 1.07E-06 |
| | | CP>NR | 2.54 | 3.11E-17 |
| L | midFrontalOrb | Intercept | 1.92 | 4.73E-09 |
| | | CP>SP | 0.91 | 0.004 |
| | | CP>NR | 1.01 | 8.90E-04 |
| L | medialFrontal | Intercept | 2.15 | 2.48E-11 |
| | | *CP>SP* | *0.49* | *0.16* |
| | | CP>NR | 1.41 | 4.49E-09 |
| L | midParietal | Intercept | 3.89 | 2.84E-14 |
| | | CP>SP | 1.28 | 2.52E-05 |
| | | CP>NR | 2.95 | 1.11E-15 |
| L | precentral_A | Intercept | 5.16 | 1.27E-12 |
| | | CP>SP | 1.03 | 9.55E-04 |
| | | CP>NR | 3.32 | 1.13E-17 |
| L | supFrontal | Intercept | 3.8 | 5.83E-18 |
| | | CP>SP | 0.79 | 0.002 |
| | | CP>NR | 2.96 | 1.14E-19 |
| L | insula | Intercept | 1.26 | 6.46E-16 |
| | | *CP>SP* | *0.28* | *0.141* |
| | | CP>NR | 0.66 | 6.33E-07 |
| R | postParietal | Intercept | 3.87 | 4.58E-18 |
| | | CP>SP | 2 | 1.09E-10 |
| | | CP>NR | 3.51 | 1.68E-19 |
| R | midFrontal | Intercept | 2.65 | 2.66E-14 |
| | | CP>SP | 1.06 | 2.62E-04 |
| | | CP>NR | 1.81 | 1.90E-09 |
| R | precentral_B | Intercept | 3.61 | 6.29E-14 |
| | | CP>SP | 1.26 | 1.61E-05 |
| | | CP>NR | 2.43 | 2.34E-13 |
| R | antParietal | Intercept | 2.12 | 5.07E-13 |
| | | CP>SP | 1.09 | 5.91E-05 |
| | | CP>NR | 1.61 | 6.99E-09 |
| R | midFrontalOrb | Intercept | 1.48 | 1.71E-05 |
| | | CP>SP | 0.87 | 0.006 |
| | | CP>NR | 0.8 | 0.016 |
| R | medialFrontal | Intercept | 1.96 | 5.46E-11 |
| | | CP>SP | 0.58 | 0.004 |

| | | | | |
|---|---|---|---|---|
| | | CP>NR | 1.09 | 2.04E-08 |
| R | midParietal | Intercept | 2.98 | 1.53E-11 |
| | | CP>SP | 1.31 | 1.14E-04 |
| | | CP>NR | 2.1 | 2.62E-09 |
| R | precentral_A | Intercept | 3.62 | 7.18E-13 |
| | | CP>SP | 1.26 | 1.38E-05 |
| | | CP>NR | 2.14 | 1.34E-11 |
| R | supFrontal | Intercept | 3.19 | 1.00E-16 |
| | | CP>SP | 1.04 | 6.87E-04 |
| | | CP>NR | 2.57 | 8.34E-14 |
| R | insula | Intercept | 1.02 | 1.12E-09 |
| | | *CP>SP* | *0.33* | *0.135* |
| | | CP>NR | 0.54 | 4.98E-04 |

***Table S2****. Responses to ScratchJr code problems (CP) vs. sentence problems (SP) and nonword reading (NR) in individual fROIs within the multiple demand system. P values are Bonferroni-corrected for the number of regions. Non-significant values are italicized and marked in gray.*

| Hemisphere | fROI | Regression Term | Beta | p value |
|---|---|---|---|---|
| L | postParietal | Intercept | 3.45 | 2.76E-14 |
| | | CP>SP | 2.13 | 1.88E-06 |
| | | CP>NR | 2.65 | 1.33E-08 |
| L | midParietal | Intercept | 2.63 | 9.39E-13 |
| | | CP>SP | 1.34 | 1.23E-04 |
| | | CP>NR | 1.65 | 2.75E-06 |
| L | antParietal | Intercept | 2.42 | 2.42E-14 |
| | | CP>SP | 1.3 | 0.001 |
| | | CP>NR | 1.42 | 2.75E-04 |
| L | supFrontal | Intercept | 2.02 | 4.13E-07 |
| | | *CP>SP* | *0.57* | *1* |
| | | CP>NR | 1.14 | 0.007 |
| L | precentral_A | Intercept | 3.17 | 1.43E-07 |
| | | *CP>SP* | *0.85* | *1* |
| | | *CP>NR* | *0.8* | *1* |
| L | precentral_B | Intercept | 3.15 | 2.98E-10 |
| | | CP>SP | 1.05 | 0.031 |
| | | CP>NR | 1.33 | 0.002 |
| L | midFrontal | Intercept | 2.61 | 2.63E-09 |
| | | CP>SP | 1.34 | 0.01 |
| | | CP>NR | 1.5 | 0.003 |
| L | midFrontalOrb | Intercept | 1.7 | 1.23E-05 |
| | | CP>SP | 1.18 | 0.017 |
| | | *CP>NR* | *0.89* | *0.182* |
| L | insula | Intercept | 1.18 | 3.61E-07 |
| | | *CP>SP* | *0.58* | *0.161* |
| | | *CP>NR* | *0.12* | *1* |
| L | medialFrontal | Intercept | 1.6 | 9.49E-07 |
| | | *CP>SP* | *0.67* | *0.416* |
| | | *CP>NR* | *0.47* | *1* |
| R | postParietal | Intercept | 3.93 | 2.11E-16 |
| | | CP>SP | 2.89 | 6.90E-07 |
| | | CP>NR | 3.22 | 6.16E-08 |
| R | midParietal | Intercept | 3.35 | 1.75E-09 |
| | | CP>SP | 2.17 | 2.12E-04 |
| | | CP>NR | 1.83 | 0.002 |

| | | | | |
|---|---|---|---|---|
| R | antParietal | Intercept | 2.4 | 1.43E-15 |
| | | CP>SP | 1.59 | 1.67E-05 |
| | | CP>NR | 1.53 | 3.58E-05 |
| R | supFrontal | Intercept | 2.32 | 1.95E-10 |
| | | CP>SP | 1.24 | 0.007 |
| | | CP>NR | 1.21 | 0.009 |
| R | precentral_A | Intercept | 3.52 | 8.22E-09 |
| | | CP>SP | 2.02 | 2.26E-06 |
| | | CP>NR | 1.27 | 0.004 |
| R | precentral_B | Intercept | 3.32 | 1.99E-10 |
| | | CP>SP | 2.15 | 1.67E-07 |
| | | CP>NR | 1.48 | 2.21E-04 |
| R | midFrontal | Intercept | 2.91 | 2.76E-11 |
| | | CP>SP | 1.77 | 4.57E-05 |
| | | CP>NR | 1.26 | 0.007 |
| R | midFrontalOrb | Intercept | 1.71 | 3.49E-06 |
| | | CP>SP | 1.26 | 0.006 |
| | | *CP>NR* | *0.34* | *1* |
| R | insula | Intercept | 1.19 | 1.02E-07 |
| | | *CP>SP* | *0.69* | *0.088* |
| | | *CP>NR* | *0.2* | *1* |
| R | medialFrontal | Intercept | 1.57 | 4.39E-08 |
| | | *CP>SP* | *0.89* | *0.064* |
| | | *CP>NR* | *0.31* | *1* |
